# Assessing base-resolution DNA mechanics on the genome scale

**DOI:** 10.1101/2022.02.16.480786

**Authors:** Wen-Jie Jiang, Congcong Hu, Xinyao Yi, Qianyi Xu, Tianai Lou, Haojie Wang, Jialu Zhou, Hanwen Zhu, Yong Li, Yilin Wen, Ruiqi Fan, Jing Shen, Catherine C.L. Wong, Xiaoqi Zhen, Hua-Jun Wu

**Affiliations:** Peking University Cancer Hospital and Institute, Beijing, China; Center for Precision Medicine Multi-Omics Research, Peking University Health Science Center, Beijing, China; School of Basic Medical Sciences, Peking University Health Science Center, Beijing, China; Central Laboratory, Key Laboratory of Carcinogenesis and Translational Research (Ministry of Education), Peking University Cancer Hospital and Institute, Beijing 100142, China; Department of Mathematics, Shanghai Normal University, Shanghai, China; University of California, San Diego; Trinity College Dublin; Peking University First Hospital, Beijing 100034, China; Peking-Tsinghua Center for Life Sciences, Beijing 100871, China; Advanced Innovation Center for Human Brain Protection, Capital Medical University, Beijing 100069, China; Department of Gynecology and Obstetrics, Chinese PLA General Hospital, Beijing 100853, China

## Abstract

Intrinsic DNA properties such as bending play a crucial role in diverse biological systems. A recent advantage in the high-throughput method called loop-seq makes it possible to determine bendability of hundred thousand 50-bp DNA duplexes in one experiment. However, it’s still infeasible to assess whole sequence bendability in large genomes such as human, which needs thousands of loop-seq experiments. Here we introduce ‘BendNet’ – a neural network to accurately predict the intrinsic DNA bending at base-resolution by only given DNA sequences. BendNet can increase the resolution of experimental results, and can predict DNA bendability for any new given sequences in high accuracy. We applied BendNet to the human genome and observed high-stiffness regions located at both transcriptional start sites and transcriptional end sites. Such stiffness patterns are different for coding and non-coding genes, which matches distinct nucleosome occupancy patterns. As expected, most transcription factors (TFs) bind in DNA of low bendability. In contrast, we observed an unusually high bendability within binding elements of specific TFs such as EBF1 and regulators of genome folding such as CTCF. These factors either co-bind or compete with nucleosomes to carry out their functions. More interestingly, CTCF binding regions exhibit the highest bendability than other DNA elements, implying their potential role in trapping and holding the CTCF in the exact locations to make sure CTCF as stable anchor in loop extrusion process. Our work provides a tool to assess DNA bendability for large-scale DNA sequences and expands our understanding on DNA mechanics in chromatin regulation and genome folding.

## Introduction

DNA-protein interactions are essential for many key cellular processes, including DNA replication^1^, chromatin formation^2^ and transcriptional regulation^3^. These interactions require DNA bending to embrace proteins, which involve intrinsic properties of DNA fragments likely below 100 base-pair (bp)^4^. Assays, such as electrophoretic mobility^5^ and single-molecule fluorescence resonance energy transfer (smFRET)^6^, have been developed to determine the DNA bending ability (termed as DNA bendability). Specifically, they measure the looping rate of a single DNA fragment of approximately 100-bp in length at a time, therefore are limited by their throughput. Recent advance in a sequencing-based approach called loop-seq ^7^,^8^ has led to vast increase in throughput, which has scaled-up looping rate detection from dozens to hundred thousand DNA duplexes in one study through combining smFRET and systematic enrichment of ligands by exponential enrichment (SELEX)^9^ selection methods. In the same study, they applied loop-seq to demonstrate the contribution of DNA bendability to nucleosome organization in yeast through measuring DNA looping rate that tile the regions of interest at 7 bp resolution^8^. However, it remains infeasible to tile larger genomes such as human at base resolution, because library preparation of this method requires synthesizing equivalent numbers of DNA duplexes to the genome size that relies on constructing thousands of such libraries, and this is not considering the sequence variations between individuals.

To extend bendability assessment to any DNA duplexes in large sequence space, we developed a deeplearning-based method termed BendNet, which extracts and learns sequence features encoding DNA bendability using capsule networks^10^ without routing^11^. BendNet can predict DNA bendability of any given sequences with high agreement to that measured by loop-seq and other low throughput approaches. We also demonstrated that BendNet can accurately provide a finer map of DNA bendability in yeast whole genome at base-resolution as compared to loop-seq study^8^, resulting in the same biological findings. To explore the utility of BendNet, we applied it to human genome and obtained the first time base-resolution map of DNA bendability in mammals. We found DNA bendability might be involved in not only transcription initiation but also transcription termination, and it displays specific patterns for coding and non-coding genes. Moreover, DNA bendability of cis-elements can determine the possibility of the transcription factors (TFs) binding and co-binding. In addition, majority of TFs show low bendability in the binding center compared to the surrounding regions, while minority of TFs display elevated bendability, with either co-bind or compete with histones. Notably, CTCF binding sites show the highest bendability peaks implying the anchor function of CTCF to form the 3D folding architecture of the genome in loop extrusion process. These findings help us to understand the DNA mechanics and its association to transcription regulation and nucleosome occupancy in mammals. Our method can also be applied to other DNA mechanics such as twisting or supercoiling when similar datasets are available.

## Results

### Overview of BendNet

We developed BendNet, inspired by capsule network^10,11^ and homogeneous vector capsules^12^, to predict bendability of DNA duplex from sequence alone (Fig. 1a). The BendNet architecture consists of a convolutional module and a capsule module (Fig.1a). The convolutional module includes multiple consecutive convolutional blocks, each of which has three stacked convolutional layers with increased kernel and channel sizes followed by dropout and batch normalization. The output of each convolutional block is fed into a capsule block, which contains a capsule, dropout and batch normalization. The results of all capsule blocks are stacked, and subsequently to a fully connected layer to output a regression score as bendability prediction.

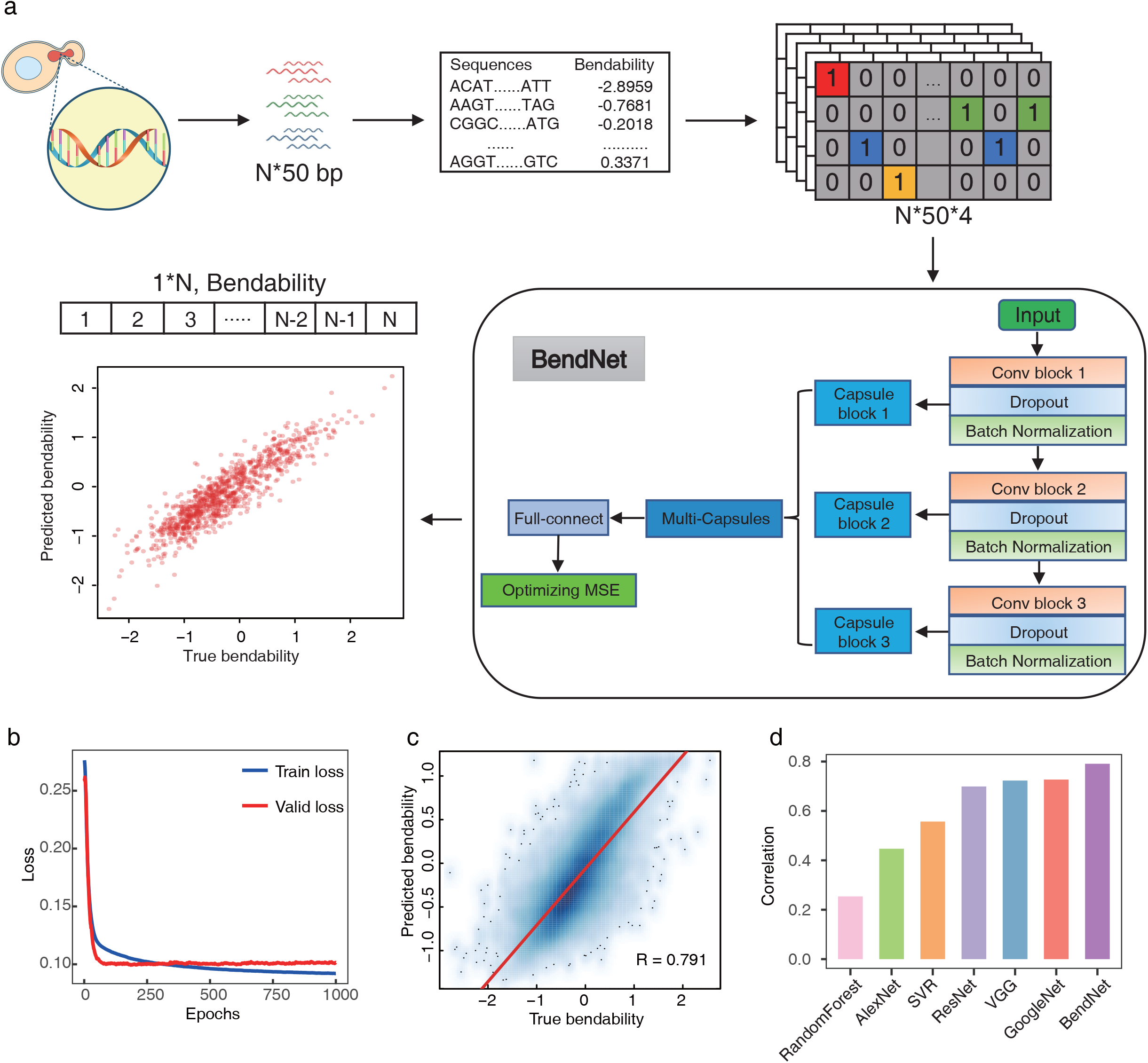

We trained BendNet on the dataset of intrinsic DNA bendability measured by loop-seq^7^. The dataset contains bendability of 270,806 DNA duplexes from five independent experiments including random, reference and codon-altered sequences in different regions of interest in yeast. We first examined the data distribution, and found an extremely high frequency at 0.02096348 (over 9000 DNA duplexes). We removed these data points, because an abnormal response score can introduce bias to the prediction model in statistical learning^13^, and obtained bendability of 264,860 DNA duplexes as DNABend dataset. We split the dataset into 70% training, 20% validation and 10% hold-out test to proceed training. BendNet learned the optimal architecture by using a genetic algorithm^14^ proposed by *Mitchell* in training and validation sets, and achieved Pearson correlation between predictions and loop-seq measurements of 0.793 in the test set. We compared this primary BendNet model to other state-of-the-art machine-learning and deep learning models including Random Forest (RF), Support Vector Regression (SVR), AlexNet, VGG, GoogleNet, and ResNet (Fig. 1b). We observed at least 5% higher correlation than other deep learning methods, and 20% higher correlation than classical machine learning methods (Fig.1c). In addition, the training cost of BendNet per epoch is at least 14 times lower than other tested deep learning models (Fig.S1 j).

To demonstrate the potential of BendNet to uncover biological discoveries, we hold out one of the five independent experimental data which contains bendability of DNA duplexes tiling the chromosome V of yeast genome in 7-bp resolution as test set, and trained another BendNet model. The model achieved Pearson correlation of 0.757 with loop-seq result, which is slightly lower than that of our primary BendNet model but still better than that of other deep learning models. A visualization of BendNet predictions and loop-seq measurements in chromosome V depicts an overall high agreement between them (Fig. 2a). In detail, the predictions of BendNet are as sensitive as loop-seq measurements in detecting DNA bendability in individual regions (Fig. 2a), and successfully uncovered the previous observation of low DNA bendability within nucleosome-depleted regions (NDRs)^8^.

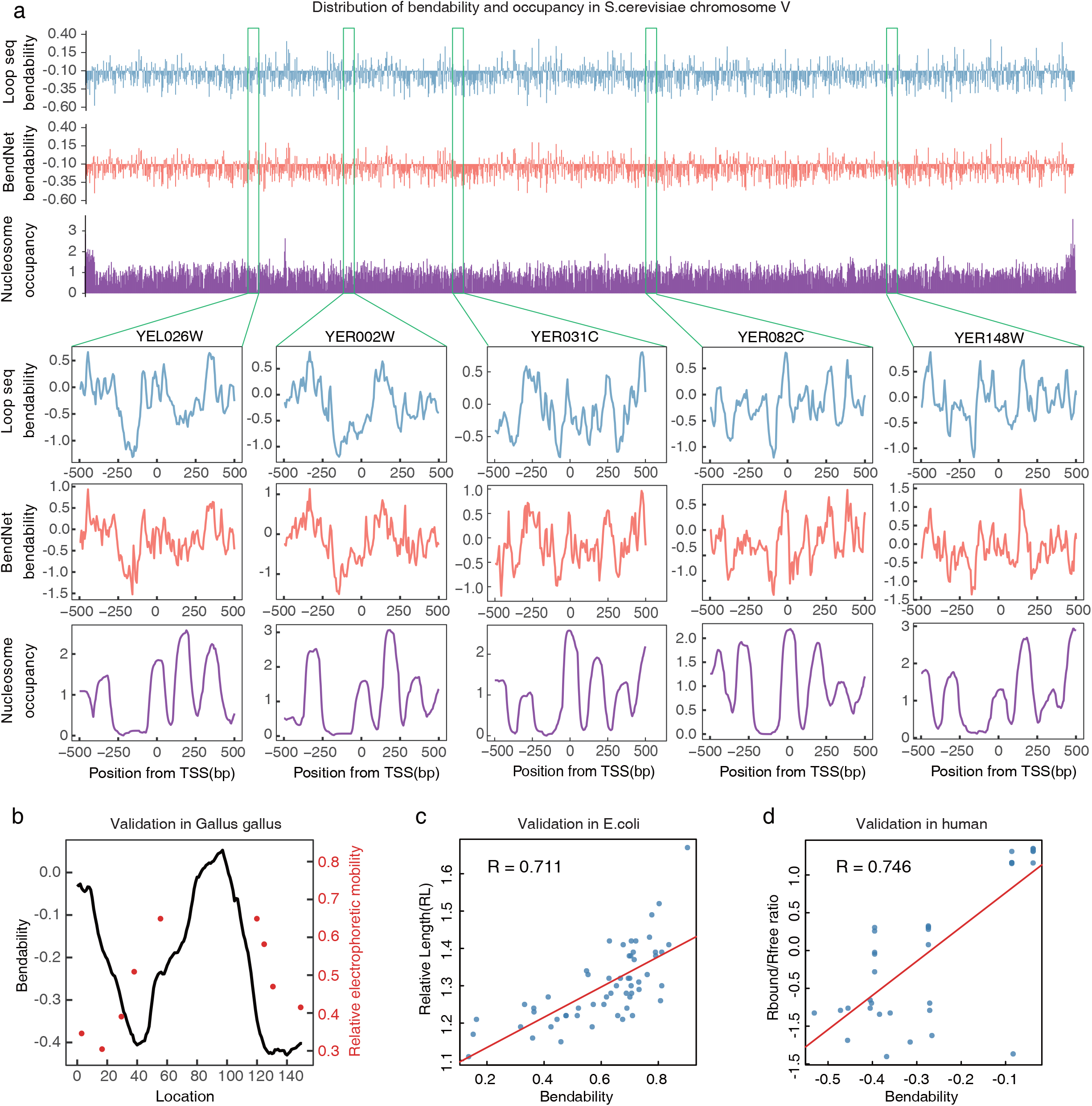

To demonstrate the generalization of BendNet, we collected the intrinsic cyclizability of DNA duplexes measured by three distinct low throughput experimental assays. The measurements include relative ionization mobilities of 9 data points of a CTCF binding region in *Gallus gallus*^15^, relative lengths (RL) of 56 DNA duplexes in *E. coli* ^16^and Rbound/Rfree ratios of 35 DNA duplexes in human ^17^. Although based on distinct protocols and performed on sequences from different species, predictions by BendNet show an overall high consistency with the above three experimental measurements (Fig. 2b-d). In particular, BendNet achieves the Pearson correlations of 0.711 in E.coli and 0.746 in the human dataset, which are comparable with the test accuracy in loop-seq experiment. These results demonstrate that BendNet is generally applicable to access DNA bendability of any given sequences across species.

### BendNet predicts base-resolution DNA bendability in human

To explore the role the DNA bendability in mammals, we applied BendNet to the human genome and obtained base-resolution DNA bendability in the whole genome scale. The bendability follows an approximately normal distribution with negative skewness (Fig.3a), which reveals an over-representation of high bendability regions likely caused by the more nucleosome-occupied regions than NDRs in the genome. A previous study discovered low bendability regions within NDRs upstream of the selected transcription start sites (TSSs) in yeast^8^. To extend our understanding on the DNA bendability in different gene types, we examined bendability in promoter regions of protein-coding gene (PCG), long non-coding RNA (lncRNA), non-coding RNA (ncRNA), and pseudogene. Generally, we observed clearly defined regions of rigid DNA (with low bendability) at both TSS (Fig. 3d) and transcription end site (TES) (Fig. 3d). More specifically, PCG exhibits the biggest bendability drop that matches with the nucleosome depletion at both TSS and TES (Fig. S2a); lncRNA shows a weaker bendability decrease than PCG, and displays a more obvious change in TES than in TSS which is coincidence with the nucleosome depletion in TES but not TSS; sncRNA demonstrates an oscillatory bendability pattern, with a notably low bendability at TES which is associated with a decreased nucleosome occupancy; pseudogene displays a minimal bendability change at both TSS and TES. These results demonstrate that the low DNA bendability is generally associated with low nucleosome occupancy at both TSS and TES, although there are distinct bendability patterns for different gene types.

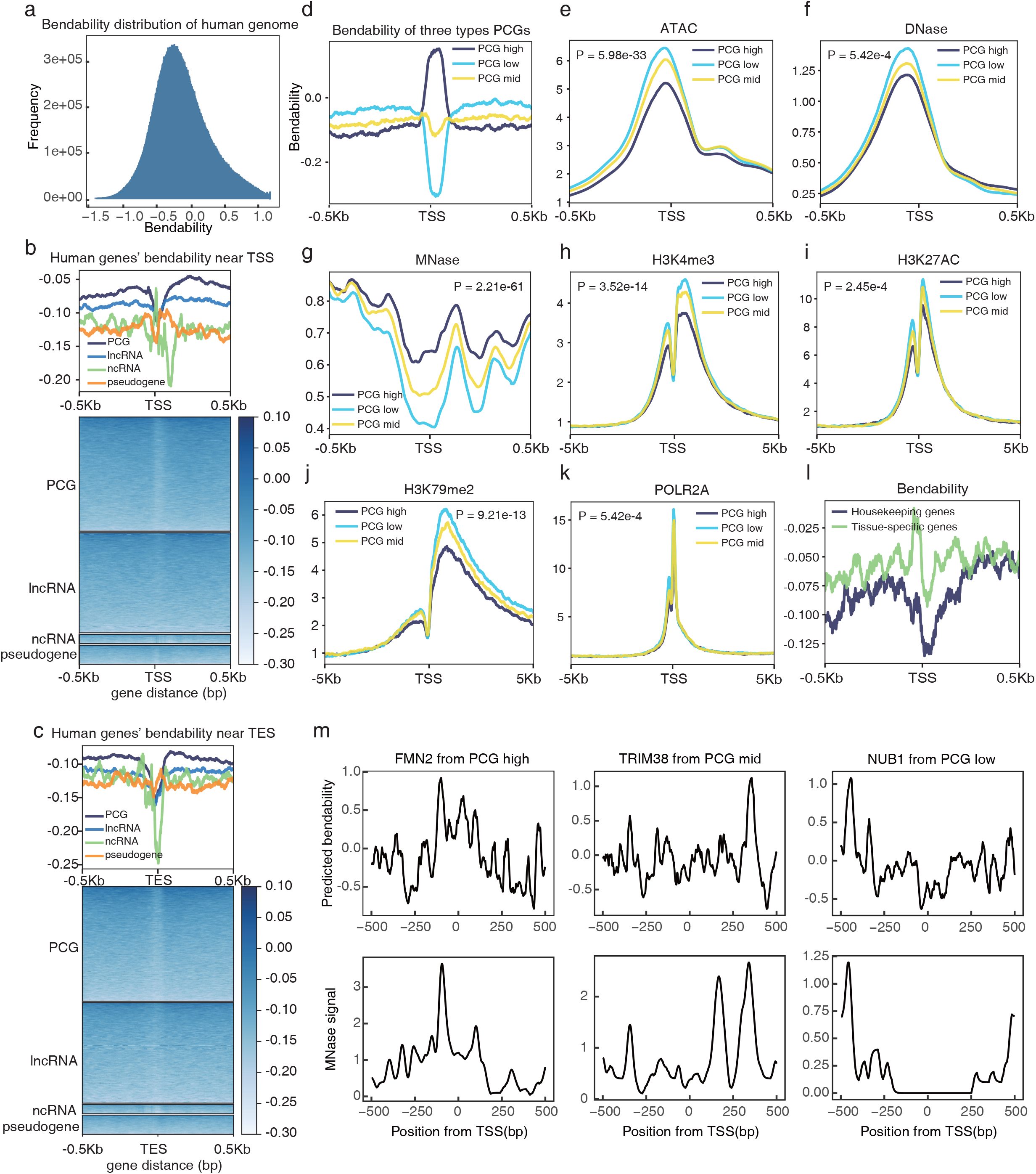

We next sought to explore the effect of DNA bendability on chromatin regulation at different gene promoters. We divided all PCGs into three equal groups (low, mid, and high) according to their average bendabilities at TSS region -20 ∼ +70 bp (Fig.3d and Fig.S2a). For each PCG in above three groups, we still observed low, mid and high bendability patterns at TSS regions, which are positively associated with nucleosome occupancy measured by MNase signal (Fig. 3m,g). By analyzing ATAC-seq and DNase-seq, we found rigid TSSs (with low bendability) are more accessible (Fig. 3e,f) than bendable TSSs (with high bendability). Moreover, the rigid TSSs are found to be enriched with the RNA polymerase II (Pol II) and H3K79me2 ChIP-seq signals, which are associated with transcriptional initiation and elongation, respectively. Consistent with this observation, the histone modifications of transcriptional activation, such as H3K4me3 and H3K27ac, are also enriched in rigid TSSs. This indicates the potential role of DNA mechanics in transcriptional regulation through chromatin regulation. In addition, we observed that the rigid TSSs are more conserved across species than bendable TSSs (Fig. S2e) and enriched for the housekeeping genes (Fig.3l), which suggests that rigid TSSs are functionally essential during both evolution and cellular development.

To explore the influence of the rigid TSS on TES, we also investigated the bendability of TES within the three groups. It is worth noting that the bendability at TSS is not associated with that at the same TES (Fig. S2d). Therefore, we performed the same analysis on TES without considering their bendability patterns at TSS, and found XXX.

### Effect of DNA bendability on TF binding

Next, we sought to investigate the effect of DNA bendability on transcription factor (TF) binding. By analyzing the ChIP-seq data of 152 TFs in GM12878, we found distinct bendability patterns in TF binding sites, which is quantified by the Pearson correlation between the average profile of bendability and ChIP-seq signal at TF binding regions (Fig. 4a). At the TF binding sites, over 2/3 of TFs such as CREB1 exhibit depressed bendability (Fig. 4b), about 1/3 of TFs such as CTCF display elevated bendability, some TFs in between such as STAT5A shows no clear bendability patterns. We then applied another quantification by measuring the bendability height in the TF binding regions, and observed the consistent patterns (Fig. 4a).

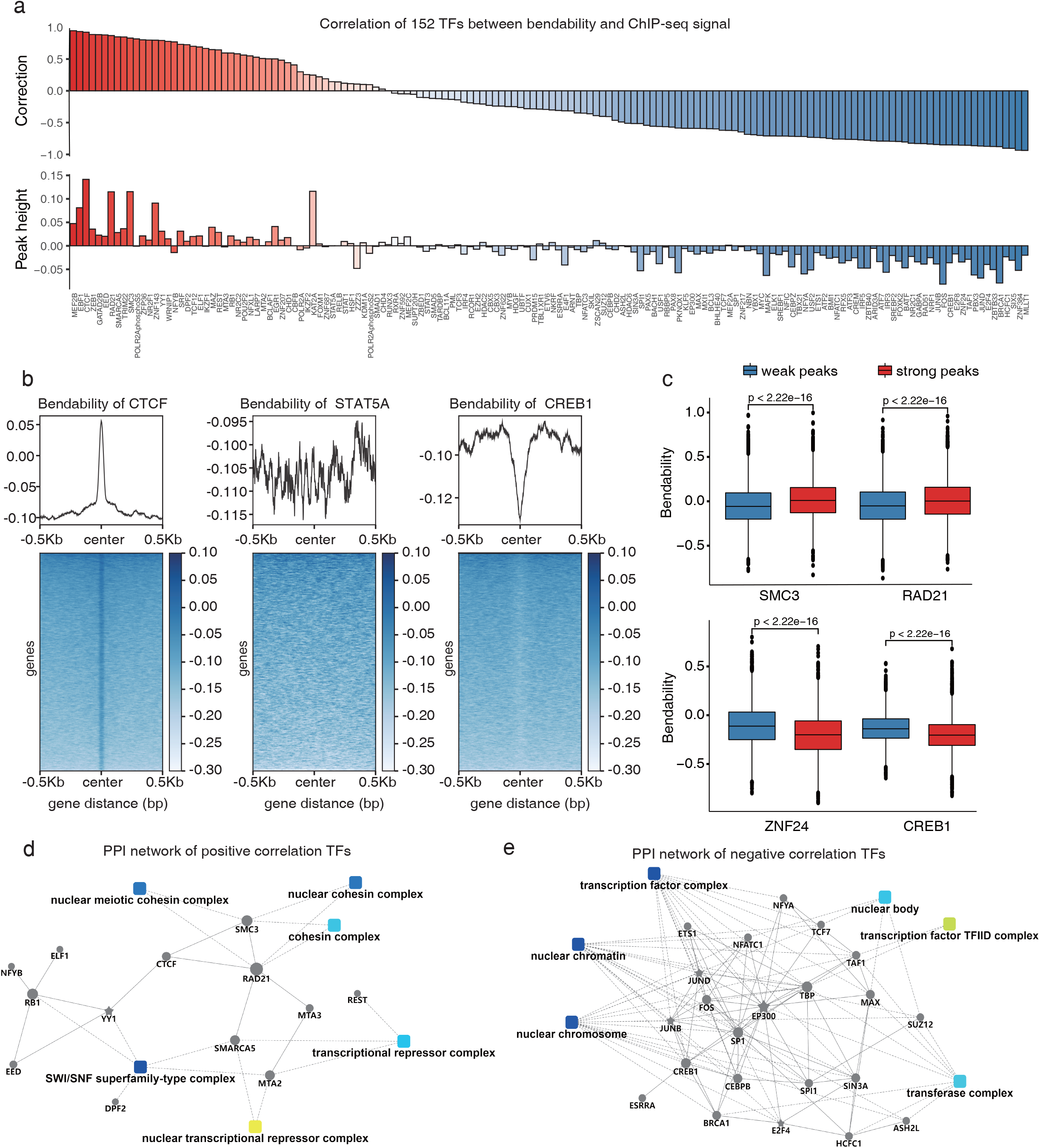

We then performed GO enrichment analysis ^18 19^ and the protein-protein interaction (PPI) ^20^ network analysis on TFs with rigid (Pearson R < -0.3) and bendable (Pearson R > 0.3) binding sites. We found that TFs with rigid binding sites are enriched for transcription factor complex, nuclear chromosome, and transferase complex (Fig.4e), while TFs with bendable binding sites are enriched for cohesin, transcriptional repressor, and SWI/SNF superfamily-type complexes (Fig.4d). Further analyses demonstrate that the transcriptional activators are enriched in TFs with overall rigid binding sites, while the transcriptional repressors are enriched in TFs with overall bendable binding sites.

To further explore the effect of bendability on the binding strength of individual TFs, we obtained binding peaks with the top and bottom quantiles of ChIP-seq signals, and investigated the bendability difference between them for each of the 152 TFs. We found strong binding peaks exhibit lower bendability than weak ones for the TFs with overall rigid binding sites such as CREB1 and ZNF24, and reversed pattern for the TFs with overall bendable binding sites such as SMC3 and RAD21 (Fig. 4c). These findings indicate the DNA mechanics plays a complex role in determining the TF binding.

### DNA bendability shapes TF binding and co-binding in promoter

Next, we sought to investigate the functional role of DNA bendability for TFs with overall rigid binding sites, which are mostly located in the gene promoter regions (Fig. 4a). Most TFs regulate gene expression through binding and co-binding on cis-regulatory elements in the promoter and enhancer regions. As is commonly known, sequence motif is critical for the TF binding but can not perfectly explain it; what other DNA properties contribute to the TF binding remains elusive. Here, we show the TF binding and co-binding are influenced by the DNA intrinsic bendability. To better illustrate it, we focused on the top 30 TFs with the highest promoter binding frequency. We found an enriched binding in the rigid TSSs for every TFs with bendability increase compared to that in the bendable TSSs (Fig. 5a). Moreover, the rigid TSSs are also enriched for the co-binding of these TFs (Fig. 5b). However, the enrichment of co-binding in rigid TSSs could be induced by the enrichment of single TF binding. To study this confounder, we calculated the co-binding ratio by adjusting for the single TF binding frequency (see Methods), and found co-binding event is still enriched in most of the rigid TSSs. These findings indicate the promoter DNA bendability is critical to both TF binding and co-binding, and is an intrinsic regulator of gene transcription other than the sequence motif.

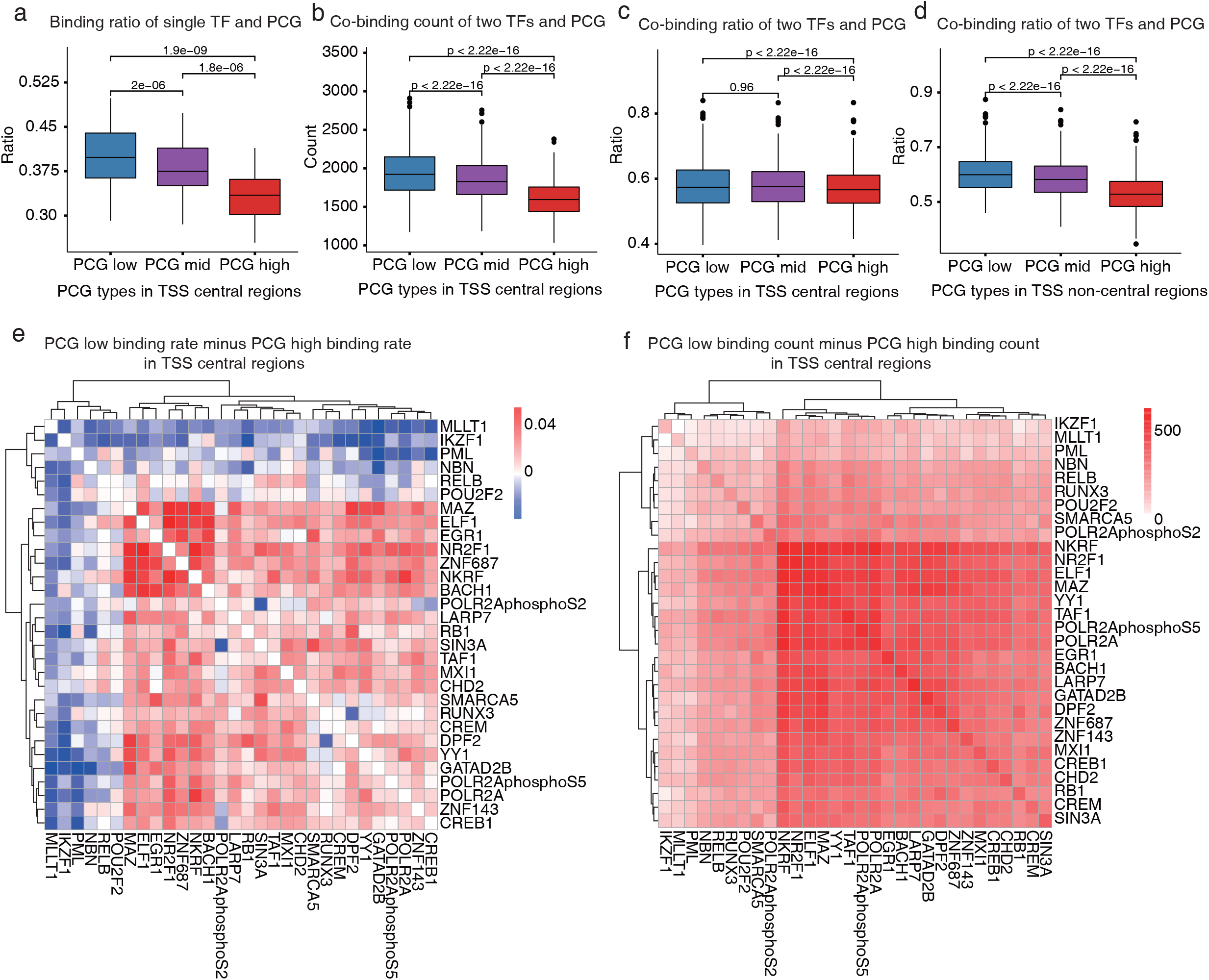

### Effect of unusually bendable DNA binding sites

We then sought to investigate the functional implication of DNA bendability for TFs with overall bendable binding sites (Fig. 4a and Fig. S3c,d). As illustrated in Figure 4d, these TFs are mostly involved in 3D genome folding, and co-bind with CTCF, including CTCF itself, cohesin subunits - RAD21 and SMC3, and regulators of promoter-enhancer loops - YY1 and ZNF143. Further analyses demonstrate CTCF binding sites (CBSs) exhibit the strongest bendability peaks (Fig. S3c), while no strong bendability pattern is observed at the binding sites of the other four factors (Fig. 6i-o). The overall bendability peak of these factors is due to the overlapping with CBSs, which emphasizes the central role of CTCF in regulating the 3D genome organization through DNA bendability. Consistant with this observation, there is a well-defined NDR at the CBS with up- and down-stream nucleosomes are aligned accordingly, but much weaker patterns for the binding sites of the other factors (Fig. S4f-i). Moreover, we observed the potential CBSs if not possess CTCF binding in specific cell lines are occupied by nucleosomes. More specifically, we investigated the MNase-seq signal in potential CBSs with or without actual CTCF binding in GM12878 cells. The latter exhibits a clear nucleosome occupancy signal, while the former doesn’t (Fig. S5). This mutually exclusive pattern is observed in the other 6 tested cell lines, indicating a competitor role of CTCF with other nucleosome proteins to bind to the specific bendable regions. This phenomenon also suggests a mechanism that the CTCF and its bendable binding elements cooperate to form the anchor points in the loop formation process because the sharply defined bendable regions in the CBSs can facilitate attracting and capturing CTCF to form accurate and stable boundaries.

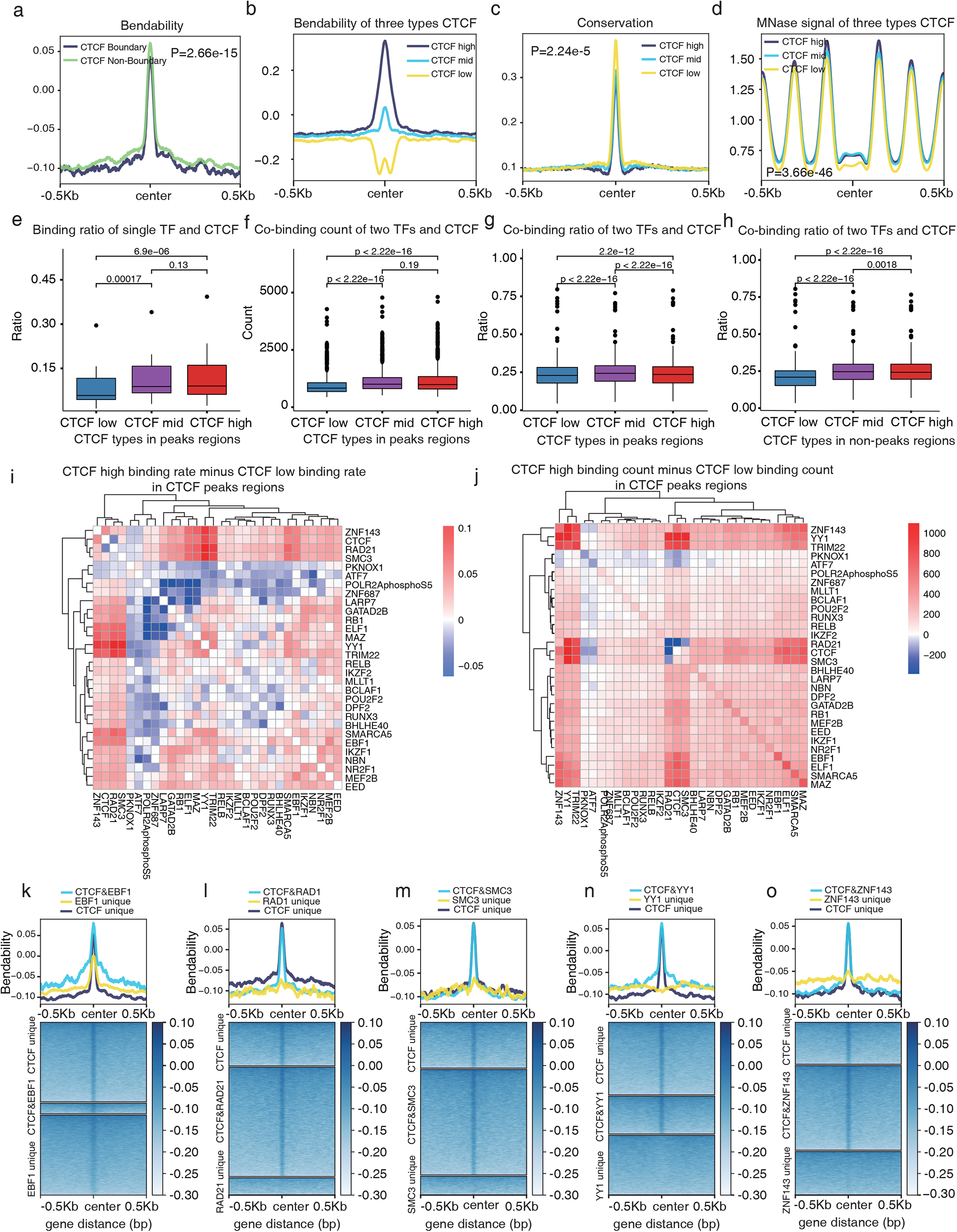

To survey the functional effect of DNA bendability in CBSs, we divided the binding sites into three equal groups (low, mid, and high) according to their average bendabilities (Fig. S3a). No obvious difference is seen in nucleosome occupancy. However, we noted higher sequence conservation across species in rigid than in bendable CBSs (Fig. 6c), which is consistent with the above results that the rigid DNA generally has higher conservation. In addition, we found the bendable CBSs are enriched for co-binding events of the other genome folding factors. These findings demonstrate a complex contribution of DNA bendability on TF co-binding, that is in promoter regions the rigid DNA contributes to enriched TF co-binding, while in CBSs the bendable DNA contributes to enriched TF co-binding. We also observed the boundary CTCF regions are enriched with bendable DNA.

Interestingly, except for the genome folding factors, there are four TFs: EBF1, EGR1, MAZ, MEF2B showing overall bendable binding sites (Fig. S3d) although it’s not as strong as CBSs. Further analyses demonstrate it is not due to the overlapping with the CBSs (Fig. 6k and Fig. S3e-g). By exploring the MNase-seq data, we found CREB1, EGR1, and MAZ are similar to CTCF, which show a well-defined NDR at their binding sites, while EBF1 binding sites exhibit a nucleosome occupied pattern. EBF1 is a pioneer factor^21^ that can directly bind DNA with nucleosome, therefore demonstrating an association between bendable DNA and nucleosome occupied pattern. These findings indicate genome folding factors, pioneer factors, and some rare TFs can have unusually bendable DNA in their binding sites that have to compete with nucleosome proteins, and functionally differently as compared to most TFs bind to the rigid DNA that intrinsically blocks the nucleosome occupancy.

## Discussion

Decoding DNA mechanics and its effect on chromatin regulation is one of the fundamental questions in genomics. Our work provides a tool to assess bendability, one of the mechanical properties of DNA, for massive-scale DNA sequences to study their biological consequences. The manuscript is, to our knowledge, the first comprehensive analysis of the only DNA mechanics in a whole genome scale in mammals. Our data provide the first base-resolution map of DNA bendability prediction in human, which expands our knowledge of how DNA mechanics influences chromatin regulation through DNA-macromolecular interactions. Our framework can be easily applied to assess other DNA mechanics, such as DNA twisting, supercoiling, and torsional rigidity when enough experimental measurement data is available, and a multi-task learning strategy could further improve the prediction accuracy. BendNet can be applied to the other genomes to study the variation of DNA bendability during evolution and its effect on gene function, and be used to investigate the effect of DNA mechanics on diverse biological systems, such as nucleosome assembly during DNA replication, genome stability maintenance, epigenetic inheritance, DNA damage repair, V(D)J recombination in B- or T-cell development.

## Method

### Training and evaluation data

Our model was trained and evaluated based on a set of DNA bendability measured by loop-seq, which consists of bendability of 270,806 DNA duplexes from five independent experiments including random, reference and codon-altered sequences in different regions of interest in *S*.*cerevisiae*. After removing the outliers with extremely high frequency and averaging the bendabilities of DNA duplexes with multiple measurements, we obtained in total 264,860 valid DNA duplexes and their bendability measurements as our processed data for the following analyses.

For training the primary model, 264,860 DNA duplexes were randomly split into 70% training (185,402), 20% validation (52,972) and 10% hold-out test (26,486) sets. For training the second model, we hold out one of the five independent experimental data which contains bendability of DNA duplexes tiling the chromosome V of yeast genome in 7-bp resolution as test set (82,404), and randomly split the left data into training (146,010) and validation sets (36,502) with splitting ratio of 8:2.

### Model architecture

The BendNet architecture consists of a convolutional module and a capsule module. The convolutional module includes multiple sequential convolutional blocks, each of which has three stacked convolutional layers with increased numbers of kernels followed by dropout and batch normalization. The output of each convolutional block is fed into a capsule block, which contains a capsule, dropout and batch normalization layers. The results of all capsule blocks are stacked, and subsequently output to a fully connected layer to produce a regression score as bendability prediction. More specifically,

1. **Input:** BendNet takes one-hot encoding of DNA sequences as input, i.e., N*50-bp DNA sequences were first transformed into N*50*4 matrices and flattern to one dimension as the input to the model. To increase the generalization of our model, we randomly reversed 50% of the input DNA sequences in training, validation and test sets.
2. **Convolutional module:** This module contains multiple convolutional blocks, each of which has three convolutional layers with kernel sizes 2*1, 3*1, and 4*1, respectively. To capture the global structure of the input sequences, we used increased number of kernels for each convolutional block. In detail, three layers of the first convolutional block have 16 kernels of size 2*1, 32 kernels of size 3*1 and 64 kernels of size 4*1, while the numbers are 32, 64, and 128 for the second convolutional block, and so on. The number of convolutional blocks is a hyperparameter to be determined by the validation set.
3. **Capsule module:** The capsule module consists of three capsule blocks, and each capsule block contains a capsule, a dropout, and a batch normalization layers.
4. **Output:** Results of three capsule blocks are stacked and output to a dense layer to get a regression score.

BendNet with three convolutional blocks contains 287,442 parameters, including 285,484 trainable parameters and 1,958 non-trainable parameters. The mean squared error (MSE) is used as the loss function, and the Adaptive Moment Estimation algorithm is used to update the parameters. The model with the minimum MSE on the validation set in 1000 epochs was used as the final model. BendNet was written in Python using the TensorFlow and Keras framework.

### Hyperparameter optimization approach

Our model has two types of hyperparameters, i.e., structural hyperparameters including the number of convolutional/capsule modules, the number of capsule classes, and the number of dimensions, and nonstructural hyperparameters including learning rate, dropout rate in convolutional and capsule blocks. We adopted different strategies to optimize two types of hyperparameters. For three structure-related hyperparameters which will increase the model complexity, we used control variates to assign different values to the three hyperparameters and recorded their minimum validation loss in 100 epochs. The hyperparameters are retained when validation loss is no longer reduced significantly with the increase of their values. In other words, if two models with different hyperparamters are comparable in accuracy, we will adopt the simple one with small hyperparameters.

The genetic algorithm is adopted to tune the nonstructural hyperparamters (ref). Basically, the validation accuracy and xxx are used to measure the fitness of a set of hyperparamters. By introducing xxxx, xxx and xx as mutation and cross operations,. We generated 50 generations in this genetic algorithm each of which was composed of 20 individuals. The best two individuals that have the minimum validation loss in 100 epochs in each generation were selected to corss and mutate to produce the next generation. The best performance was achieved with a learning rate of 0.02585, dropout rate of 0.17632 in convolutional block, and dropout rate of 0.14818 in capsule block.

### Model comparison

We compared BendNet with two widely used machine learning models and four state-of-the-art deep learning models on the same dataset.

1. **RandomForest:** Random Forest is an algorithm that uses multiple decision trees to train, classify and predict samples.
2. **SVR**: Support Vector Machines (SVM) is a binary classification model whose basic model is a linear classifier defined in feature space with the largest interval, distinguishing it from the perceptron model. SVM also includes nuclear tricks, which makes it a substantially nonlinear classifier. The learning strategy of SVM is interval maximization, and its application in regression is called SVR.
3. **AlexNet:** AlexNet is the 2012 ImageNet competition winner designed by Hinton and his student Alex Krizhevsky. It has a total of eight layers, initially used for classification problems, and by changing its output layer and loss function it can be used for regression problems.
4. **VGG:**The VGG model was the second winner of the ILSVRC competition in 2014 and performed well in multiple transfer learning tasks. The 16-layer VGG was used for model comparison.
5. **GoogleNet:** GoogleNet is a deep neural network model based on the inception module launched by Google. It won the ImageNet competition in 2014. In our study, we used a classic 22-layer network.
6. **ResNet:** ResNet was ImageNet’s champion in 2015 and surpassed human recognition accuracy for the first time, reducing the error rate to 3.57%. It solved the problem of gradient explosion and gradient disappearance in deep neural network learning. ResNet with 34 layers was used in the comparison.

### Validation datasets by other experimental technologies

1. ***Gallus gallus***: This dataset includes the relative electrophoretic mobilities of 7 DNA fragments in CTCF binding regions of Gallus gallus. In detail, DNA fragments are loaded into pBEND2 plasmid, and xxxx. The relative electrophoretic mobility is a measure indicating whether the corresponding region is bendable.
2. ***E. coli***: The dataset includes relative length (RL) of xxx DNA fragments of length 57bp. According to xxx (ref), The mutations *of E. coli* DNA fragments were induced in vitro within a 57 bp region which is located at the ilvIH operon TSS -83bp∼-140bp, then 56 fragments were obtained with no redundancy. RL is defined as the ratio of the apparent length to the actual length, which is in proportion to the bendability of DNA fragment.
3. ***Human***. The dataset contains Rbound/Rfree ratios of the 35 fragments in xxx. five types of 20bp p53 response elements (p53 consensus sequence, p21/waf1/cip1, symmetric sequence, ribosomal gene cluster sequence (RGC), SV40 promoter sequence), and five recombinant plasmids (pCon30, pWaf30, pSS30,pRGC30, and pSV30) in human. These plasmids contain the above five types of p53 binding sites flanked by tandemly duplicated DNA sequences. The plasmids were cleaved at the seven restriction sites and 35 DNA fragments were obtained. The Rbound/Rfree ratio was calculated by p53DBD-DNA complex’ electrophoretic mobility compared to free DNA. A fragment with large Rbound/Rfree ratio is supposed to be more bendable.

### Annotation of human genomic

According to the GRCh38.p13 annotation for the human genome from Genecode, protein-coding gene(PCG), long non-coding RNA(lncRNA), non-coding RNA(ncRNA), and pseudogene were selected in the study.

### Classification in PCG, lncRNA, ncRNA, and pseudogene

For PCG, We divided all PCGs into three equal groups(low, mid, and high) according to their average bendability at TSS region -20∼+70bp (Fig.3 d and Fig.S2 a). Then, lncRNA, ncRNA, and pseudogene also followed this approach and were divided into three equal groups.

### Association between bendability with Chromatin opening, nucleosome modification, and transcription

We obtained the 7 kinds of sequencing such as ATAC-seq (ENCFF603BJO), DNase-seq (ENCFF960FMM), MNase-seq(ENCFF000VME), H3K4me3(<u>ENCSR057BWO</u>), H3K27ac (ENCSR000AKC), H3K79me2(<u>ENCSR000AOW</u>) and POLR2A(ENCFF328MMS) in GM12878 from ENCODE. For PCG, 3 groups(PCG low, PCG mid, and PCG high) corresponding 7 kinds of sequencing signal were obtained and test the significance between PCG low corresponding sequencing signal and PCG high corresponding sequencing signal.

### Human tissue-specific and housekeeping genes

Housekeeping genes and Tissue-specific genes were obtained from the previous study[ref].

### Association between bendability with ChIP-seq signals

We obtained the BigWig files and BED (peak locations) files of 152 TFs in GM12878 from ENCODE. We used bwtools to extract the ChIP-seq signal and bendability in the peak center ±500 bp regions of each TF.

1. **Correlation:** We took the mean value of bendability and ChIP-seq signal by column, changing the dimension from N*1000 into 1*1000. Then we calculated the Pearson correlation coefficient between ChIP-seq signal and bendability.
2. **Bendability-height**: We calculated the mean value of bendability in 1000bp regions firstly. Then the average of 200bp(±100bp) bendability in the center subtracted the average of 800bp(−500bp∼-100bp and 100bp∼500bp) bendability to get height for 152 TFs.
3. **Δbendability:** For each TF, the average value of the ChIP-seq signal corresponding to each peak was calculated, and the top 25% and the bottom 25% peaks were selected. Then the average value of bendability corresponding to each peak in the two groups(top 25% and the bottom 25% peaks) was calculated. Finally, the mean values of the two groups were subtracted to obtain bendability diff, and Wilcox was used to test the significance of the difference between the two groups.

### Protein interaction network and enrichment analysis

The protein-protein interactions were obtained from string (https://string-db.org/). The enrichment in cell composition of GO database was analyzed.

### TSS ±500 bp regions and CTCF peaks overlap with 152 TFs’ peaks

We obtained the location of TSS of all protein-coding genes and 152 TFs’ peaks, then calculated the overlap of each TF in the TSS ±500bp regions. “1” indicates overlap, “0” indicates no overlap.

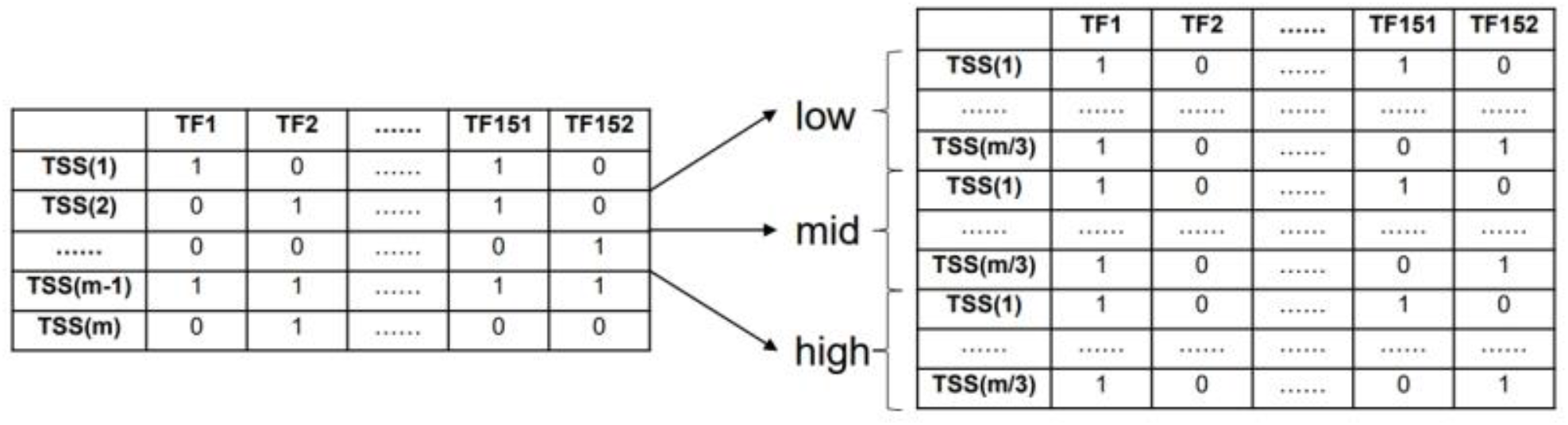

According to the average value of bendability in TSS -20bp∼+70bp regions for every gene, TSS is divided into three types (low, mid, and high) in descending order. Finally, the TSS of each category (low, mid, and high) calculated the count of 152 TFs combining transcription start regions. For example, when two TFs overlap to the same transcription start region at the same time, it is recorded as “1”, otherwise “0”, then the values of all transcription start regions were summed. Therefore we can obtain a number to evaluate the co-binding to transcription start regions of two TFs.

Since the number of peaks of each TFs is not the same, we should consider the actual situation of each TFs during normalization. sum(TF1∩TF2)/sum(TF1∪TF2) was used to calculate the degree of co-binding between two TFs, TF1 means the vector of TF1 binding to TSS, TF2 means the vector of TF2 binding to TSS.

The above was the “TSS peaks” calculation method. “TSS base” used the mean value of bendability of TSS base region(−500bp∼-20bp and 70bp∼500bp) to divide into three types (low, mid, and high) from low to high, and the subsequent calculation process is the same as TSS peak. “CTCF Peak” classification like “TSS peaks” used the average value of bendability of ChIP-seq peaks instead of a fixed region, while “CTCF base” used the average value of bendability of 1000bp region excluding ChIP-seq peak for classification “TSS peaks”.

### Sequence conservation

Sequence conservation information was obtained from the UCSC “phastCons46wayPlacental” track.

### Difference of CBS cross cell lines

We obtain MNase-seq and CTCF ChIP-seq of 6 cell lines, such as Bcell([bigwig](https://www.ncbi.nlm.nih.gov/geo/query/acc.cgi?acc=GSM1278008), [bed](https://www.encodeproject.org/files/ENCFF390ICZ/)), GM12878(<u>bigwig</u>, [bed](https://www.encodeproject.org/files/ENCFF138REW/)), H1([bigwig](https://www.ncbi.nlm.nih.gov/geo/query/acc.cgi?acc=GSM1194220), [bed](https://www.encodeproject.org/files/ENCFF023LAA/)), H9([bigwig](https://www.ncbi.nlm.nih.gov/geo/query/acc.cgi?acc=GSM1194221), [bed](https://www.encodeproject.org/files/ENCFF691YKL/)), Hela([bigwig](https://www.ncbi.nlm.nih.gov/geo/query/acc.cgi?acc=GSM2680344), [bed](https://www.encodeproject.org/files/ENCFF111RWV/)), and K562([bigwig](https://www.encodeproject.org/files/ENCFF000VNN/), [bed](https://www.encodeproject.org/files/ENCFF660GHM/)). Firstly, we merge CTCF peaks of 6 cell lines, then use bwtools to select common peaks and uncommon peaks for every cell line. Finally, calculate the average bendability and MNase signal for unique peaks and common peaks in 6 cell lines.

### Data visualization

Heatmaps and average plots were generated by deeptools. The other plots were drawn in the R environment using basic plot functions, ggplot2, and Pheatmap packages.

## Data availability

The data that supports this study are available from The corresponding authors upon reasonable request.

Saccharomyces cerevisiae genome: [Saccharomyces cerevisiae](http://ftp.ensembl.org/pub/release-105/fasta/saccharomyces_cerevisiae/dna/Saccharomyces_cerevisiae.R64-1-1.dna_sm.toplevel.fa.gz)

Human genome: [GRCh38.p13](https://www.encodeproject.org/files/ENCFF660GHM/)

Human genome annotation: [GRCh38.p13 GTF](https://ftp.ebi.ac.uk/pub/databases/gencode/Gencode_human/release_39/gencode.v39.chr_patch_hapl_scaff.annotation.gtf.gz)

ATAC-seq(ENCFF603BJO), DNase-seq(ENCFF960FMM), MNase-seq(ENCFF000VME), H3K4me3(<u>ENCSR057BWO</u>), H3K27ac(ENCSR000AKC), H3K79me2(<u>ENCSR000AOW</u>) and POLR2A(ENCFF328MMS) of GM12878 from Encode.

152TFs ChIP-seq bigwig and bed files of GM12878 were listed in the following.

ARID3A(ENCFF531CLQ, ENCFF830CXC); ARNT(ENCFF596RIT, ENCFF343TRG); ASH2L(ENCFF552EPA, ENCFF384BWK); ATF2(ENCFF973FKS, ENCFF329LQR); ATF3(ENCFF882AEU, ENCFF243XPP); ATF7(ENCFF277FJJ, ENCFF751UNU); BACH1(ENCFF631RZH, ENCFF900OJJ); BATF(ENCFF728KFD, ENCFF013ZCI); BCL11A(ENCFF824QXX, ENCFF925SGV); BCL3(ENCFF452NTN, ENCFF546WTN); BCLAF1(ENCFF185BRL, ENCFF225WFX); BHLHE40(ENCFF633YUQ, ENCFF206VRV); BMI1(ENCFF160PIE, ENCFF128CNB); BRCA1(ENCFF303GOQ, ENCFF017KXT); CBFB(ENCFF876CUT, ENCFF670LHO); CBX3(ENCFF670JSE, ENCFF196UXL); CBX5(ENCFF864AYJ, ENCFF771TRB); CEBPB(ENCFF955YFB, ENCFF019EWO); CEBPZ(ENCFF639KLW, ENCFF420SKF); CHD1(ENCFF753HYW, ENCFF143QTK); CHD2(ENCFF351VXQ, ENCFF138PNE); CHD4(ENCFF403JIP, ENCFF118QVK); CREB1(ENCFF053UZX, ENCFF795MDW); CREM(ENCFF294IGP, ENCFF657GQY); CTCF(ENCFF138REW, ENCFF485CGE); CUX1(ENCFF451AII, ENCFF745BDD); DPF2(ENCFF003ZLG, ENCFF197LVJ); E2F4(ENCFF744QAC, ENCFF482TVC); E2F8(ENCFF961ZFW, ENCFF364XKI); E4F1(ENCFF855PXT, ENCFF188UFJ); EBF1(ENCFF895MHN, ENCFF696PUH); EED(ENCFF519PDZ, ENCFF231WWY); EGR1(ENCFF637UJN, ENCFF750YNG); ELF1(ENCFF146SYU, ENCFF262CCB); ELK1(ENCFF164MPE, ENCFF746TCL); EP300(ENCFF718TWQ, ENCFF482JMC); ESRRA(ENCFF660PIB, ENCFF091TIJ); ETS1(ENCFF568AZT, ENCFF065YMX); ETV6(ENCFF151UJT, ENCFF894HHJ); EZH2(ENCFF339YTO, ENCFF256BYP); FOS(ENCFF571DGT, ENCFF503MRU); FOXK2(ENCFF639XYN, ENCFF626ILI); FOXM1(ENCFF549GKZ, ENCFF891YKE); GABPA(ENCFF093KLR, ENCFF959CKU); GATAD2B(ENCFF648CHL, ENCFF056XZR); HCFC1(ENCFF479QQB, ENCFF887BNK); HDAC2(ENCFF581FMI, ENCFF509PQH); HDAC6(ENCFF248JAL, ENCFF714MSZ); HDGF(ENCFF626TYH, ENCFF378BJI); HSF1(ENCFF663ODL, ENCFF428HGS); IKZF1(ENCFF819VMH, ENCFF875ABM); IKZF2(ENCFF723ZRN, ENCFF816IYS); IRF3(ENCFF785MSW, ENCFF806XAD); IRF4(ENCFF113VGD, ENCFF486IOQ); IRF5(ENCFF478SRO, ENCFF686LMA); JUNB(ENCFF912OPT, ENCFF245XUS); JUND(ENCFF023QWA, ENCFF078RFY); KAT2A(ENCFF586XDT, ENCFF306PGJ); KDM1A(ENCFF996NUD, ENCFF489BWY); KLF5(ENCFF992IBT, ENCFF190VIG); LARP7(ENCFF927IGC, ENCFF941WFR); MAFK(ENCFF436SJS, ENCFF949YIL); MAX(ENCFF361EVH, ENCFF232BTG); MAZ(ENCFF467VSF, ENCFF420YPZ); MEF2A(ENCFF826GQU, ENCFF778ICR); MEF2B(ENCFF884QQW, ENCFF931IBK); MEF2C(ENCFF238UKB, ENCFF907FVJ); MLLT1(ENCFF652GOE, ENCFF282NSB); MTA2(ENCFF957SRJ, ENCFF754MQM); MTA3(ENCFF475GKK, ENCFF544EYZ); MXI1(ENCFF376AEL, ENCFF122CNK); MYB(ENCFF705CGM, ENCFF870DIP); MYC(ENCFF214XPD, ENCFF405YWN); NBN(ENCFF511QTQ, ENCFF995DXI); NFATC1(ENCFF172KBM, ENCFF397GIR); NFATC3(ENCFF002XEC, ENCFF309YZY); NFIC(ENCFF980SDE, ENCFF766JTP); NFXL1(ENCFF057NHJ, ENCFF394DFE); NFYA(ENCFF937QTP, ENCFF499VHE); NFYB(ENCFF156MUM, ENCFF992FDL); NKRF(ENCFF794DRR, ENCFF959MJR); NR2C1(ENCFF626EEU, ENCFF675EPS); NR2C2(ENCFF782KRV, NR2C2, ENCFF835AYP); NR2F1(ENCFF591BVB, ENCFF553SLZ); NRF1(ENCFF390OQK, ENCFF902IBW); PAX5(ENCFF192WNJ, ENCFF397VPJ); PAX8(ENCFF934NVD, ENCFF759BRU); PBX3(ENCFF402DQD, ENCFF386ZNA); PKNOX1(ENCFF946JFA, ENCFF234CAE); PML(ENCFF242KVU, ENCFF044IIN); POLR2A(ENCFF886PSD, ENCFF328MMS); POLR2AphosphoS2(ENCFF643YBS, ENCFF353DID); POLR2AphosphoS5(ENCFF107UMF, ENCFF559YIO); POU2F2(ENCFF934JFA, ENCFF366GEF); PRDM15(ENCFF157HQD, ENCFF222OUW); RAD21(ENCFF834GOT, ENCFF571ZJJ); RAD51(ENCFF345VPL, ENCFF622TBK); RB1(ENCFF844GNV, ENCFF860LHP); RBBP5(ENCFF333IML, ENCFF707OFR); RCOR1(ENCFF874HBT, ENCFF972XNQ); RELB(ENCFF968MBN, ENCFF099LLQ); REST(ENCFF262MRD, ENCFF845YET); RFX5(ENCFF601WHE, ENCFF536CFB); RUNX3(ENCFF346JDW, ENCFF298OKU); RXRA(ENCFF063KLF, ENCFF870ANL); SIN3A(ENCFF013QNV, ENCFF858OPR); SIX5(ENCFF878FBM, ENCFF308TWB); SKIL(ENCFF506SOD, ENCFF292AVI); SMAD1(ENCFF033MGN, ENCFF596FDZ); SMAD5(ENCFF485GWN, ENCFF165TMM); SMARCA5(ENCFF550FDV, ENCFF461PLB); SMC3(ENCFF837YJA, ENCFF401YFJ); SP1(ENCFF038AVV, ENCFF952WGS); SPI1(ENCFF492ZRZ, ENCFF042MFJ); SREBF1(ENCFF397YRO, ENCFF104OMJ); SREBF2(ENCFF896MPE, ENCFF619LTN); SRF(ENCFF114CWH, ENCFF683VIM); STAT1(ENCFF689KXN, ENCFF116UVP); STAT3(ENCFF029PAA, ENCFF011FHF); STAT5A(ENCFF006GSY, ENCFF615YAM); SUPT20H(ENCFF006GSY, ENCFF141LPJ); SUZ12(ENCFF894BBO, ENCFF416XSK); TAF1(ENCFF502IPK, ENCFF307JTI); TARDBP(ENCFF847AAV, ENCFF926QFA); TBL1XR1(ENCFF276QRH, ENCFF964GST); TBP(ENCFF099PYR, ENCFF017ZEX); TBX21(ENCFF170IZO, ENCFF031SFE); TCF12(ENCFF068YYR, ENCFF579VGX); TCF3(ENCFF700TAS, ENCFF560MLE); TCF7(ENCFF900GTF, ENCFF439FNI); TRIM22(ENCFF519BGX, ENCFF980WYP); UBTF(ENCFF014PHS, ENCFF397EAZ); USF1(ENCFF879TPT, ENCFF029MAS); USF2(ENCFF196JQW, ENCFF899OET); WRNIP1(ENCFF855UKX, ENCFF956PLX); YBX1(ENCFF488IGF, ENCFF624SLC); YY1(ENCFF446ZSX, ENCFF408AOH); ZBED1(ENCFF753UAX, ENCFF758XVK); ZBTB33(ENCFF917XFN, ENCFF544CDL); ZBTB40(ENCFF171FMR, ENCFF421NUI); ZEB1(ENCFF703XCL, ENCFF807OAU); ZFP36(ENCFF627JQX, ENCFF810RXW); ZNF143(ENCFF012FCL, ENCFF191EGO); ZNF207(ENCFF854NOE, ENCFF274XTP); ZNF217(ENCFF131ALE, ENCFF341CIW); ZNF24(ENCFF664TYB, ENCFF873IIC); ZNF384(ENCFF955FRU, ENCFF862AYM); ZNF592(ENCFF510ZBJ, ENCFF574SCI); ZNF622(ENCFF359PUY, ENCFF145ZUK); ZNF687(ENCFF734BGL, ENCFF371AND); ZSCAN29(ENCFF828NYA, ENCFF843PHT); ZZZ3(ENCFF535WUH, ENCFF676HVZ);

---

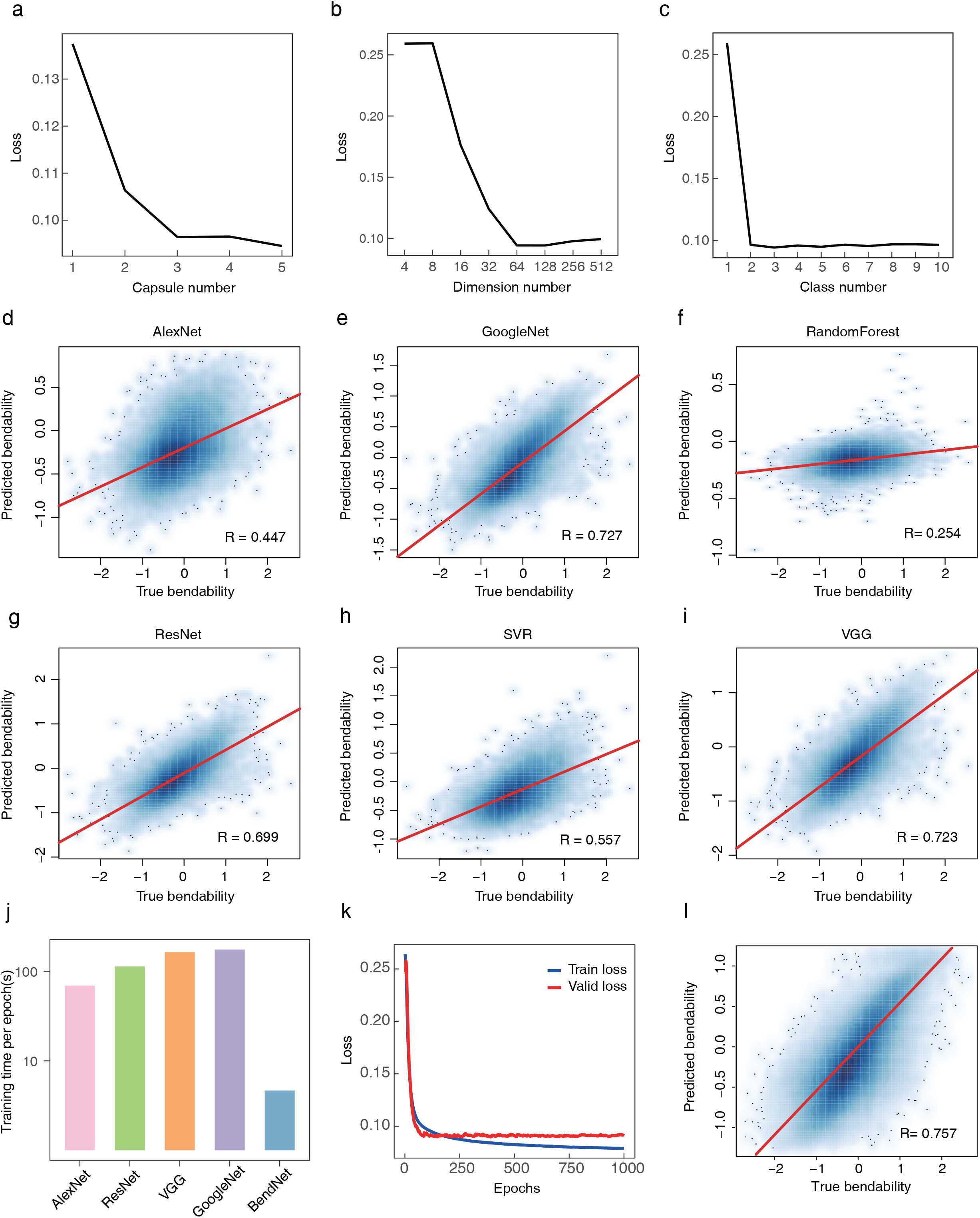

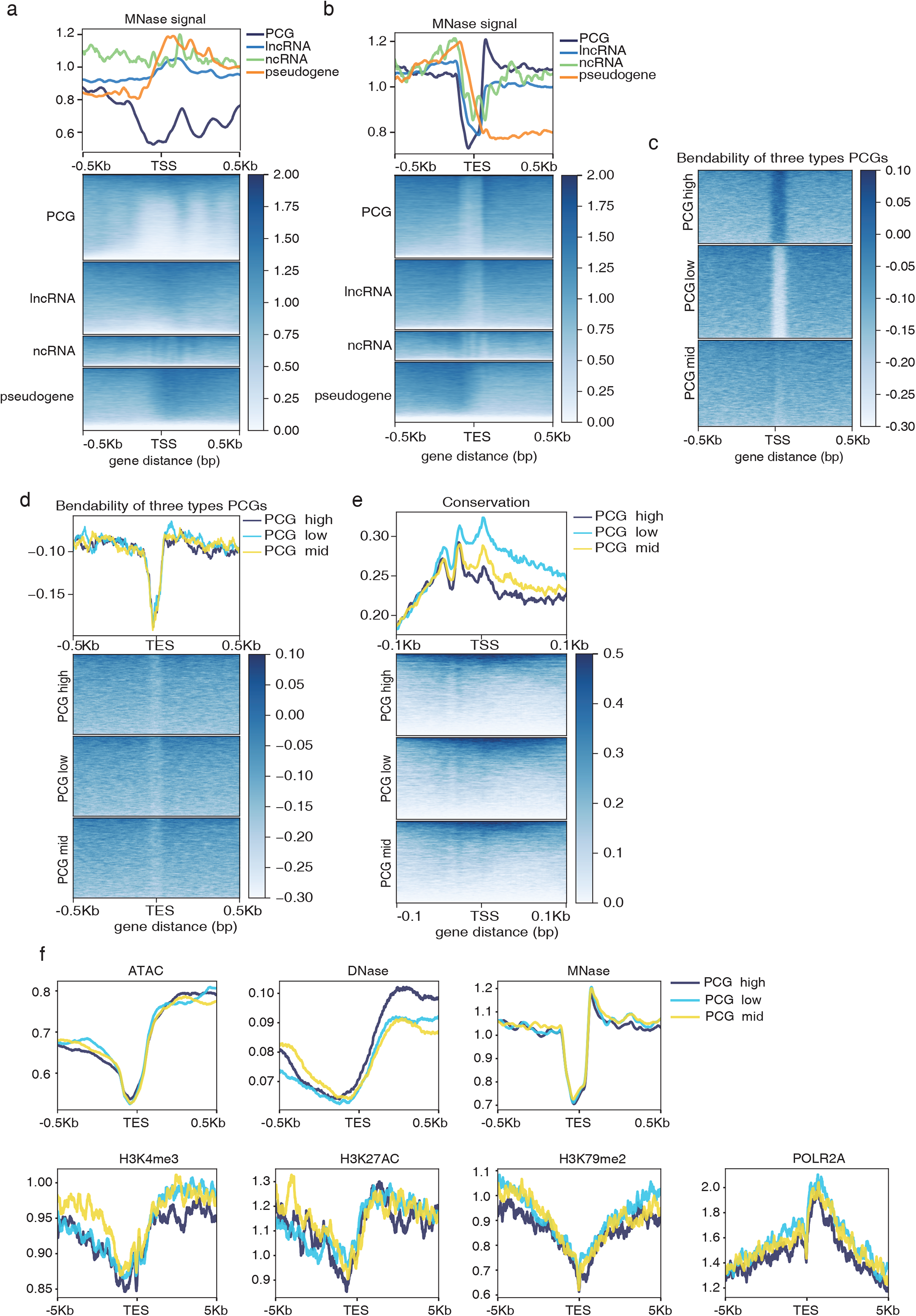

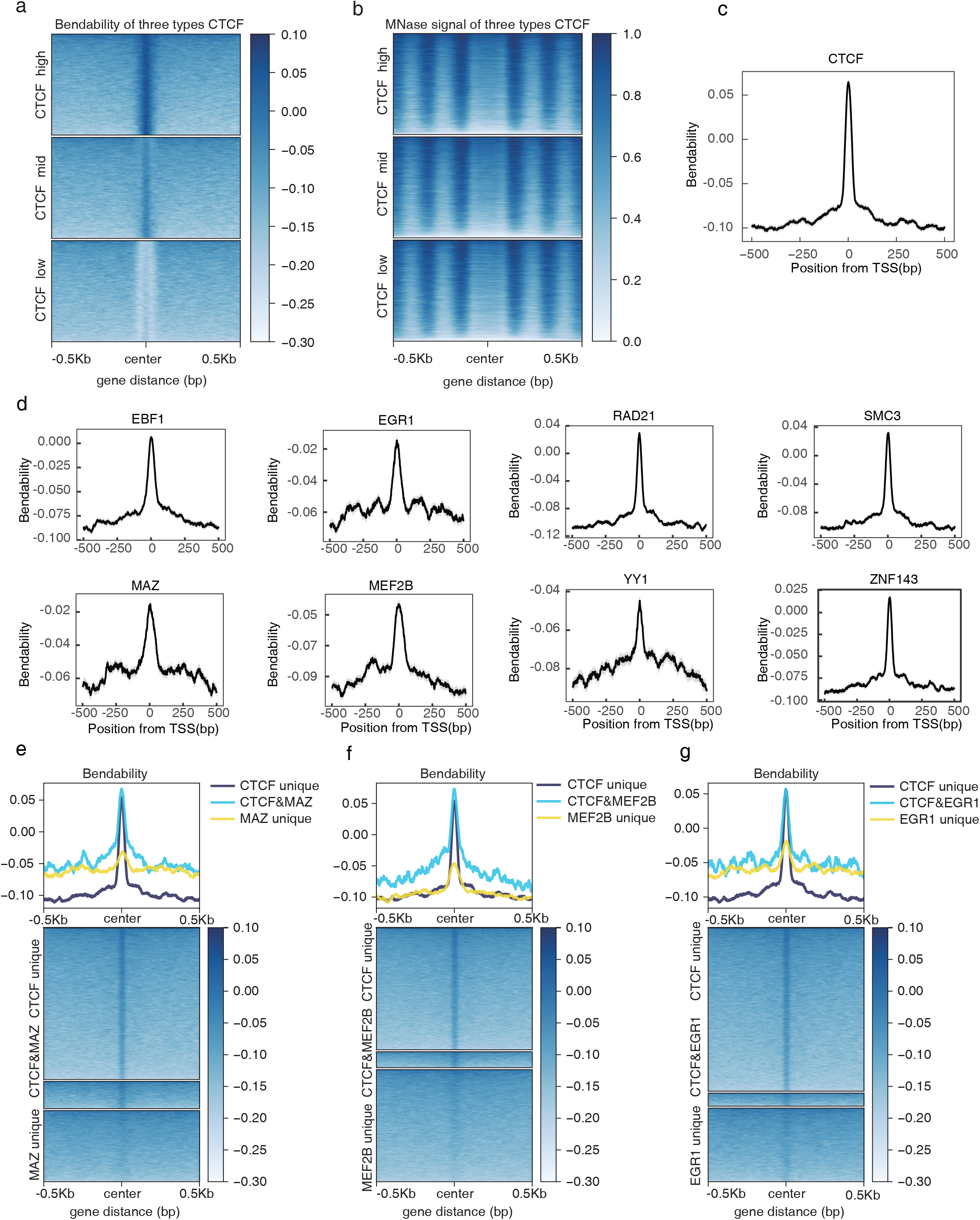

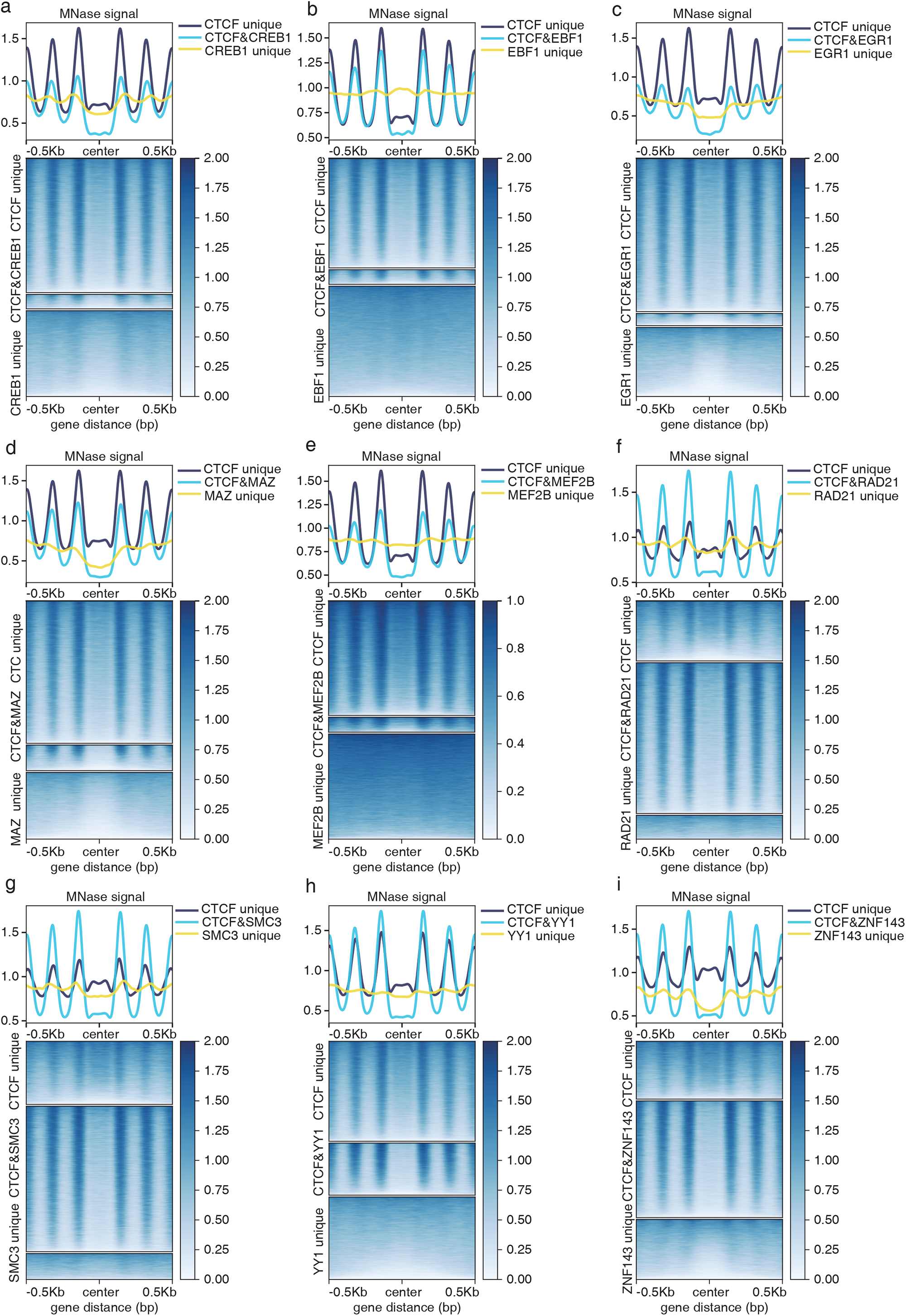

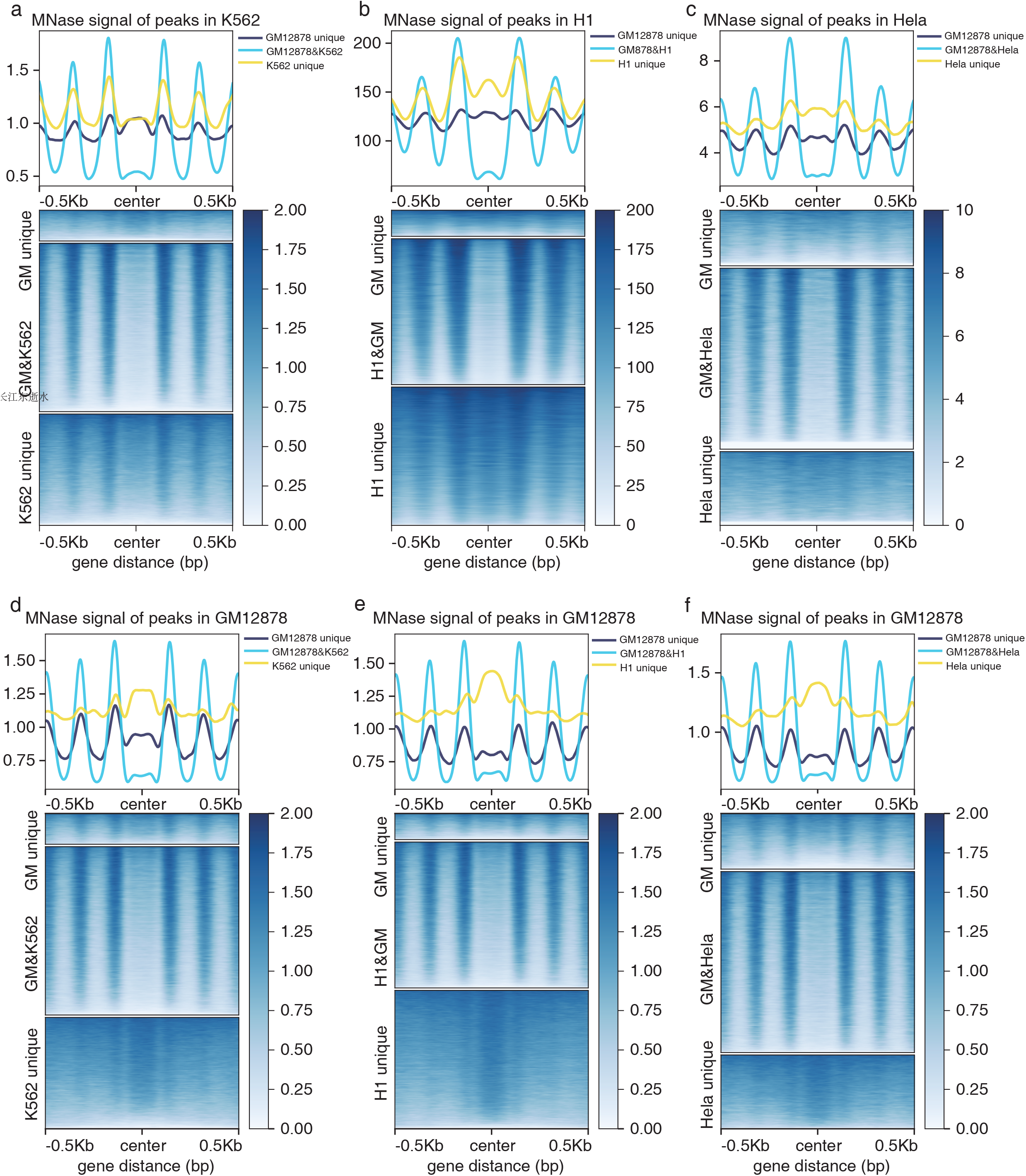

