## Supplementary material for "Assessing base-resolution DNA mechanics on the genome scale": Figure5

Fig.5: **The binding strength of top 30 TFs in TSS regions**

**a,** Binding rate of the top 30 TFs which bind to three types of PCGs in Fig.3d in TSS ±500bp regions**.** Top 30 TFs were picked by those binding sites who have the highest overlapping rate with PCGs’ TSS ±500bp regions (diagonal of Fig. S7**d,e,f**). **b,** Co-binding rate of two different TFs bind to three types PCGs in Fig.3d synchronously in the TSS ±500bp region (upper triangle except diagonal of Fig.S7**d,e,f**). **c,** Co-binding ratio of three types PCGs in Fig.3d bind to two different TFs synchronously in the TSS ± 500bp region (upper triangle except diagonal of Fig.S7**a,b,c**). **d,** Co-binding ratio of three types PCGs (PCG low, PCG mid, and PCG high) bind to two different TFs synchronously in the TSS ± 500bp region. PCGs were divided into three types by bendability in TSS non-central regions, i.e. TSS -500bp to -20bp and +70bp to +500bp regions also called ‘PCG low’, ‘PCG mid’, and ‘PCG high’. **e,** Deviation of binding ratio between two types PCGs (PCG low and PCG high in Fig.3d) binds to two different TFs synchronously in TSS ±500bp regions (Fig.S7 **a,c**). **f.** Deviation of binding count between two types PCGs (PCG low and PCG high in Fig.3d) binds to two different TFs synchronously in the TSS ±500bp regions (Fig.S7 **d,f**).
