## Supplementary material for "Assessing base-resolution DNA mechanics on the genome scale": Figure6

Fig.6: **The binding strength of top 30 TFs and CTCF**

**a,** Bendability of CTCF boundary and non-boundary. The P value came from *wilcox* *test*. **b,** CTCF was classified into three categories (CTCF low, CTCF mid, and CTCF high) by the average bendability of CTCF peaks regions from low to high. The plot profiles three types of CTCF in CTCF peaks central ±500bp regions. **c,d,** Profile of conservation (c), and MNase signal (d) corresponding to CTCF categories in Fig.6b. **e,** Binding rate of the top 30 TFs which bind to the CTCF peaks central ±500bp regions of the three types CTCF in Fig.6b. Top 30 TFs were picked whose binding sites have the highest overlapping rate with CTCF peaks central ±500bp regions (diagonal of Fig.S8 **d,e,f**). **f,** Co-binding rate of two different TFs bind to three types CTCF in Fig.6b synchronously in the CTCF peaks central ±500bp regions (upper triangle except diagonal of Fig.S8 **d,e,f**). **g,** Co-binding ratio of three types CTCF in Fig.6b binds to two different TFs synchronously in the CTCF peaks central ±500bp regions (upper triangle except diagonal of Fig.S8**a,b,c**). **h,** Co-binding ratio of three types CTCF (CTCF low, CTCF mid, and CTCF high) binds to two different TFs synchronously in the CTCF peaks central ±500bp regions. CTCF were divided into three types by bendability in CTCF non-peaks’ regions, i.e. CTCF peaks central ±500bp regions except peaks’ regions called ‘CTCF low’, ‘CTCF mid’, and ‘CTCF high’. **i,** Deviation of binding ratio between two types CTCF (CTCF low and CTCF high in Fig.6b) binds to two different TFs synchronously in CTCF peaks central ±500bp regions (Fig.S8 **a,c**). **j.** Deviation of binding count between two types CTCF (CTCF low and CTCF high in Fig.6b) binds to two different TFs synchronously in CTCF peaks central ±500bp regions (Fig.S8 **d,f**). **k-o,** The profiles of CTCF unique peaks (dark blue), CTCF peaks co-bind to EBF1 **(k)**, RAD21 **(l)**, SMC3 **(m)**, YY1 **(n)**, and ZNF143 **(o)** (sky blue), and EBF1 unique peaks **(k)**, RAD21 unique peaks **(l)**, SMC3 unique peaks **(m)**, YY1 unique peaks **(n)** and ZNF143 unique peaks **(o)** (yellow).
