## Supplementary material for "Assessing base-resolution DNA mechanics on the genome scale": FigureS1

Fig.S1: **Model’s optimization and comparison.**

**a,b,c,** Parameters optimization of BendNet structure, including capsule number **(a)**, dimension number **(b)**, and class number **(c)**. **d-i,** Correlations between bendability predicted by six models and true bendability in Dataset1, the color gradient shows the density of the dots. **j,** Average training time for RF, AlexNet, SVR, ResNet, VGG, GoogleNet, and BendNet per epoch. **k,** Training and validation loss of 1000 epochs in Dataset2. **l,** Correlation of true bendability and predicted bendability in Dateset2, the color gradient shows the density of the dots.
