## Supplementary material for "Assessing base-resolution DNA mechanics on the genome scale": FigureS2

Fig.S2: **Profiles of PCGs around TSS or TES regions**

**a,b** The line plot and heatmap of MNase signal (y axis) in the TSS ±500bp region (x axis) and TES ±500bp region (x axis) of four typical human genes (PCG, lncRNA, ncRNA, and pseudogene). PCG, protein-coding genes**.** lncRNA, long non-coding RNA. ncRNA, non-coding RNA. **c,** Profiles of bendability corresponding to the three types PCGs (in Fig.3d) in TSS ±500bp regions. **d,** Bendability profiles of three types of PCGs (in Fig.3d) in TES ±500bp regions. **e**, Profiles of conservation corresponding to the three types of PCGs (in Fig.3d) in TSS ±500bp regions. **f,** Profile of ATAC, DNase, MNase, H3K4me3, H3K27AC, H3K79me2, and POLR2A associated with three types PCGs (in Fig.3d) in TES ±500bp regions.
