## Supplementary material for "Assessing base-resolution DNA mechanics on the genome scale": FigureS3

Fig.S3: **Relationship between CTCF and other TFs in bendability**

**a,** Heatmap of three types CTCF in Fig.6b. **b,** MNase-seq signal corresponding to three types CTCF in Fig.6b. **c,** Bendability of CTCF. **d,** Bendability of EBF1, EGR1, RAD21, SMC3, MAZ, MEF2B, YY1, and ZNF143. **e-g,** Profiles of CTCF unique peaks (dark blue), CTCF peaks co-bind to MAZ **(e)**, MEF2B **(f)**, and EGR1 **(g)** (sky blue), and MAZ unique peaks **(e)**, MEF2B unique peaks **(f)**, and EGR1 unique peaks **(g)** (yellow).
