## Supplementary material for "Assessing base-resolution DNA mechanics on the genome scale": FigureS4

Fig.S11: **MNase-seq signal of peaks between CTCF and other TFs**

**a-i,** MNase signal of CTCF unique peaks (dark blue), CTCF peaks co-bind to CREB1 peaks **(a)**, EBF1 peaks **(b)**, EGR1 peaks **(c)**, MAZ peaks **(d)**, MEF2B peaks **(e)**, RAD21 peaks **(f)**, SMC3 peaks **(g)**, YY1 peaks **(h)**, ZNF143 peaks **(i)** (sky blue), and CREB1 unique peaks **(a)**, EBF1 unique peaks **(b),** EGR1 unique peaks **(c)**, MAZ unique peaks **(d)** MEF2B unique peaks **(e)**, RAD21 unique peaks **(f)**, SMC3 unique peaks **(g)**, YY1 unique peaks **(h)** and ZNF143 unique peaks **(i)** (yellow).
