## Supplementary material for "Assessing base-resolution DNA mechanics on the genome scale": FigureS5

Fig.S12: **MNase-seq signal of peaks between GM12878 and K562、H1、Hela cell lines**

**a,d,** MNase signal of GM12878 unique peaks (dark blue), peaks shared by GM12878 and K562 (sky blue), and K562 unique peaks (yellow) in GM12878 cell line (a) and K562 cell line (d), respectively. GM12878 refers to lymphoblastoid cell line. K562 refers to immortalized myelogenous leukemia cell line. **b, e**, MNase signal of GM12878 unique peaks (dark blue), peaks shared by GM12878 and H1 (sky blue), and H1 unique peaks (yellow) in GM12878 cell line (b) and H1 cell line (e), respectively. H1 refers to human embryonic stem cell lines. **c, f,** MNase signal of GM12878 unique peaks (dark blue), peaks shared by GM12878 and Hela (sky blue), and Hela unique peaks (yellow) in GM12878 cell line (c) and Hela cell line (f), respectively. Hela refers to human cancer cell line.
