## Supplementary material for "Assessing base-resolution DNA mechanics on the genome scale": Figure1

Fig.1: **BendNet, a multi-capsules model to predict DNA sequence’s bendability in *S.cerevisiae***

**a, C**onstructure of BendNet. We first got S.cerevisiae’s DNA fragment data with intrinsic cyclizability by “loop-seq” and transformed it to a one-hot array, followed by multiple consecutive convolutional blocks, each of which has three stacked convolutional layers with increased kernel and channel sizes followed by dropout and batch normalization. The output of each convolutional block acts as an input to construct a capsule block, which contains a capsule, dropout and batch normalization (See Method for details). The results of all capsule blocks are stacked, and followed by a fully connected layer to output a regression score as bendability prediction. **b,** Training and validation loss of 1000 epochs in Dataset1. **c,** Scatter plot of true bendability and predicted bendability in Dataset1, the color gradient indicates the density of the dots. **d,** The correlation between true bendability and predicted bendability in RandomForest, AlexNet, SVR (Support Vector Regression), ResNet, VGG, GoogleNet, and BendNet.
