## Supplementary material for "Assessing base-resolution DNA mechanics on the genome scale": Figure2

Fig.2: **Bendability’ verification in *S.cerevisiae* chromosome V, Gallus gallus*, E.coli* and Human**

**a,** Bendability (by Loop-seq and BendNet), nucleosome occupancy (top) of *S.cerevisiae* chromosome V and five individual genes (YEL026W, YER002W, YER006W, YER082C, and YER148W) among it (bottom). **b,** The line plot and scatter plot between bendability predicted by BendNet (y axis in the left), relative electrophoretic mobility (y axis in the right) and DNA sequence (x axis) in Gallus gallus. **c**, The correlation between relative length (RL) (y axis) and bendability predicted by BendNet (x axis) in *E. coli.* **d***,* The correlation between Rbound/Rfree ratio (y axis) and bendability predicted by BendNet (x axis) in Human**.**
