## Supplementary material for "Assessing base-resolution DNA mechanics on the genome scale": Figure3

Fig.3: **Prediction of bendability in the human genome**

**a,** Bendability distribution of the whole human genome. **b,c,** The line plot and heatmap of bendability (y axis) of four typical human genes (PCG, lncRNA, ncRNA, and pseudogene) in TSS (b) and TES (c) ±500bp region (x axis). TSS, transcription start site. TES, transcription end site. PCG, protein-coding genes. lncRNA, long non-coding RNA. ncRNA, non-coding RNA. **d,** PCG was classified into three categories (PCG low, PCG mid, and PCG high) by average bendability of TSS central regions, i.e. -20bp to +70bp regions from low to high. The plot shows profiles of three types PCGs in TSS ±500bp regions. **e-k,** The sequencing signal of ATAC **(e)**, DNase **(f)**, MNase **(g)**, H3K4me3 **(h)**, H3K27AC **(i)**, H3K79me2 **(j)**, and POLR2A **(k)** in TSS ± 500bp region, each of which corresponded to three types of PCGs in Fig.3d. The P-value was obtained from *Wilcox test* between average sequencing signal of PCG low and PCG high. **l,** Bendability of housekeeping genes and tissue-specific genes in TSS ±500bp region. **m,** Bendability (top) and MNase signal (bottom) in TSS ±500bp region of three individual genes FMN2、TRIM38 and NUB1 picked from three types of PCGs in Fig.3d respectively.
