## Supplementary material for "Assessing base-resolution DNA mechanics on the genome scale": Figure4

Fig.4: **The relationship between ChIP-seq signal and bendability**

**a,** Correlation of bendability and ChIP-seq signal of 152 TFs. We got GM12878 152 TFs ChIP-seq data from ENCODE (Encyclopedia of DNA elements). After that, the average value of bendability and ChIP-seq signal was calculated in each TF peak central ±500bp region. Then we obtained the 152 TFs’ correlation (Top) between bendability and ChIP-seq signal. “Bendability height” equals to the average bendability of the peak central ±100bp region minus to other regions (-500bp to -100bp and +100bp to +500bp) (bottom). TF, transcription factor. **b,** Profiles of three different TFs’ (CTCF, STAT5A and CREB1) bendability whose correlations were 0.9213, 0.1436, -0.8507 in Fig.4a respectively. **c.** The boxplot of bendability for weak peaks and strong peaks of four individual TFs. “weak peaks” and “strong peaks” correspond to 25% lowest and 25% highest ChIP-seq peaks after sorting the peaks' average ChIP-seq signal. **d,e,** PPI network of positive **(d)** and negative **(e)** correlation (|R|>0.3) between ChIP-seq signal and bendability.
